# Tracking antigen-specific T cell response to cancer immunotherapy

**DOI:** 10.1101/2024.11.29.626027

**Authors:** I.A. Shagina, T.O. Nakonechanaya, A.V. Izosimova, D.V. Yuzhakova, V.D. Skatova, K.R. Lupyr, M. Izraelson, A.N. Davydov, M. Shugay, O.V. Britanova, D.M. Chudakov, G.V. Sharonov

## Abstract

In each human or model animal, T cell receptor (TCR) of each effector/memory T cell clone recognizes from one to several cognate peptide-MHC complexes (pMHC). Limited knowledge on TCR repertoire specificities restricts our capacity to rationally interpret this information, both diagnostically and in preclinical research. Here we 1) develop cost-efficient wet lab and computational pipeline to identify mouse TCRs specific to particular peptides, 2) produce dataset of helper T cells (Th) TCR beta CDR3s specific to B16 melanoma neoantigens in the I-Ab pMHCII context, available in VDJdb, and 3) apply this dataset to track tumor-specific T cell response to the CTLA4 blocking immunotherapy on orthotopic B16 melanoma model. We show that tumor promotes clonally independent Tregs carrying tumor-specific TCRs characteristic for Th cells. We further show that CTLA4 blockade promotes Th clonal expansion and induces general, non-tumor-specific Th-to-Treg plasticity. Altogether, we provide a universal pipeline for the investigation of mouse T cell responses at the antigen-specific level, facilitating development and validation of immunotherapeutic and vaccination approaches.

## Introduction

Immunotherapeutic approaches based on checkpoints inhibition profoundly advanced cancer treatment over the past decades, and this progress continues, now empowered by efficient anti-cancer vaccines arising (1,2). The aim of immunotherapeutic intervention is to disrupt or counteract tumor-mediated immunosuppression (3–5) and to provoke efficient clonal T cell and NK cell antitumor cytotoxic response.

In preclinical models, we principally should be able to reproducibly track the abundance, tissue localization, and expansion of tumor-specific T cell clones in diverse immunotherapeutic and vaccination settings, based on the bulk TCR repertoire profiling (TCR-Seq) or scRNA-Seq. However, the usability of this attractive option is currently hampered by our limited knowledge of particular tumor antigens-specific TCRs.

Here we develop a medium-throughput, cost efficient, vaccination-based pipeline that allows rapid identification of public mice TCRβ CDR3 motifs recognizing particular peptide antigens by means of comparative TCR-Seq data analysis between vaccinated mice groups. The obtained data can be further employed for TCR repertoire-informed development and validation of immunotherapeutic and vaccination approaches to fighting cancers and infections.

We apply this pipeline to a set of previously reported immunogenic B16F10 melanoma antigens (6,7) to discover 59 TCRβ CDR3 motifs and 485 TCRβ CDR3 amino acid variants of C57BL/6 helper T cells recognizing 14 different peptides (**Fig. 1a**). The obtained dataset was uploaded to the VDJdb database (8–10) to fuel TCR repertoire-instructed investigation of immunotherapeutic approaches and vaccination strategies.

**Figure 1.**
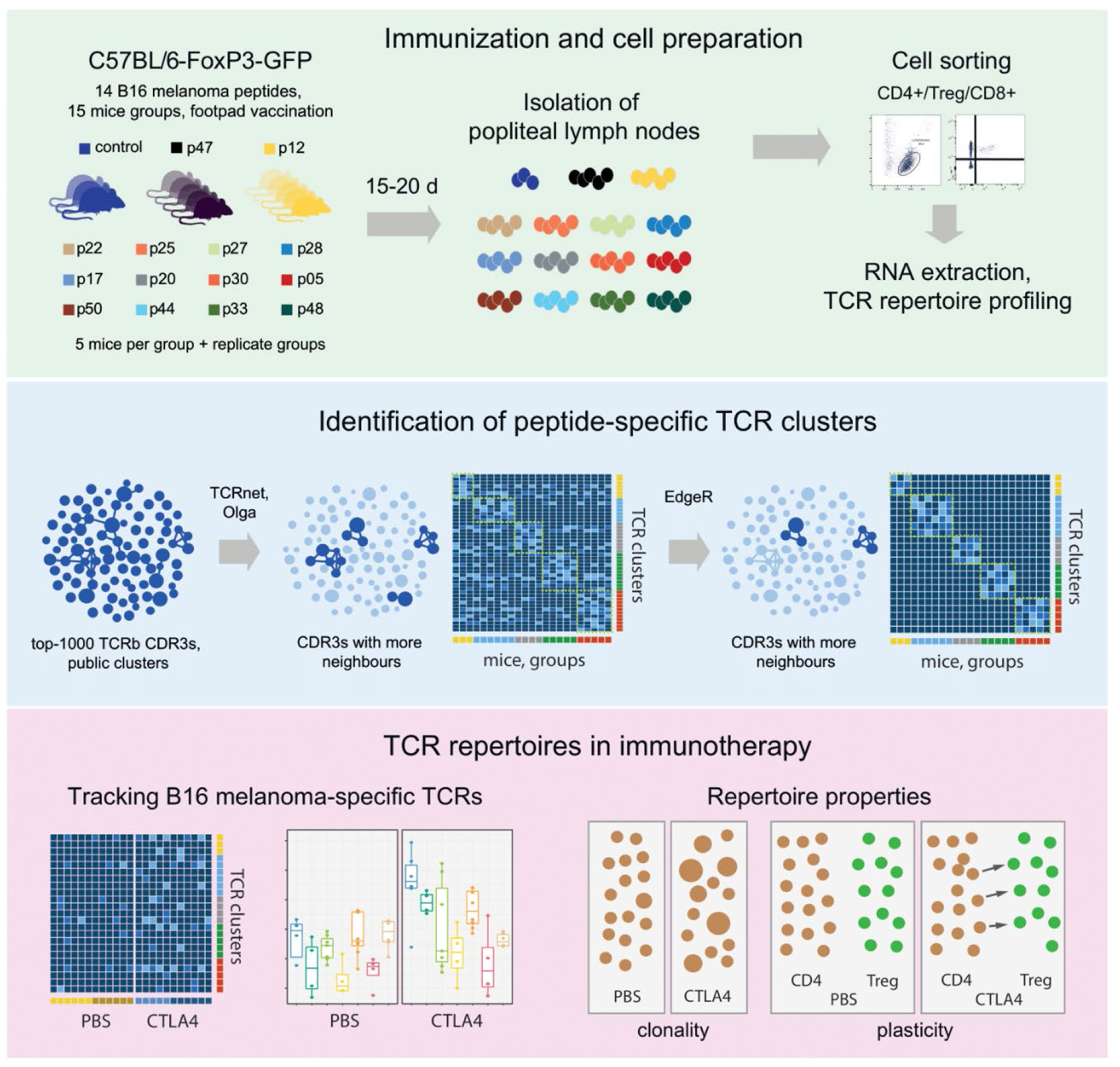
Pipeline overview. Upper panel: peptide vaccination. Middle panel: getting public antigen-specific TCR clusters. Bottom panel: TCR-informed tracking and investigation of tumor-specific T cell response.

As an illustrative example, we used TCRβ CDR3 repertoires to track B16-specific T cell response to the CTLA4-blocking immunotherapy of mice inoculated with B16F0 melanoma. We show that presence of tumor promotes *de novo* generation of Treg clones carrying tumor-specific TCRs characteristic for Th cells. Used CDR3 sequences are similar yet not identical, indicating the independent priming of convergent Th and Treg clones (11), but not Th to Treg plasticity as a driving force of this phenomenon.

As for the Th to Treg plasticity, at the same time, we show that suppression of CTLA4-mediated Treg function promotes Th clones conversion to Treg phenotype, but this process is not specifically targeted at the tumor-specific Th clones.

## Materials and Methods

### Peptides and vaccination

Neoepitope peptides were synthesized by GenScript with >85% purity. According to the solubility testing provided by the manufacturer, peptides p05, p20 and p30 were dissolved in DMSO at 40 mg/ml and others were dissolved in mQ water at 20 mg/ml. Peptide stock solutions were aliquoted and stored at −80°C for long term and −20°C for short term (3 month). For vaccination, peptides were diluted in PBS (Sigma-Aldrich) at 400 μg/ml and thoroughly mixed with incomplete Freund’s adjuvant (IFA) at 1:1 ratio by 1 ml syringe.

The experiments were carried out on transgenic C57BL/6-FoxP3^EGFP^ mouse strain (kindly provided by Alexander Rudensky, Sloan Kettering Institute, New York, NY, USA). This strain was initially generated by knocking in the eGPF gene subcloned into the first exon of FoxP3 gene. Homozygous female FoxP3-EGFP^+/+^ mice were selected by PCR-typing so that all Tregs were EGFP+.

3–6-month-old female mice were anesthetized with intramuscular injection (i.m.) of 100 μg Zoletil 100 (Virbac Sante Animale). 100 μl of peptide IFA mix containing 20 μg of peptide was injected subcutaneously (s.c.) in each hindfoot.

2-3 weeks after vaccination mice were euthanized with Isofluran (Esteve Pharmaceuticals, Milan, Italy). Popliteal LNs were isolated, dissected thoroughly with scissors and incubated for 30 min at 37°C in 417 µg/ml Liberase TL solution (Roche) supplemented with 10 µg/ml DNAse I (Roche). Then suspension was passed through a 70 µm cell strainer (BD Biosciences), washed twice with Hank’s solution and stained for FACS sorting.

### Tumors and anti-CTLA4 therapy

A B16F0 murine melanoma cell line was used in the study. B16F0 cells were cultured in DMEM medium (Thermo) supplemented with 10% FBS (Gibco), 0,06% L-glutamine (Paneco), 50 units/mL penicillin and 50 μg/mL streptomycin (both from Paneco, Russia) in 25mm cell flasks (Corning). After thaw cells were cultured in a CO2-incubator at 37°C and 5% CO2 for one week. For inoculation cells were trypsinized, washed twice in PBS, counted and resuspended at 1×10^6^ in 3 ml PBS.

8-11 weeks old male C57BL/6-FoxP3^EGFP^ mice were used in these experiments. Mice were anesthetized with i.m. injection of 100 μg Zoletil 100 and right flanks were shaved for better tumor control. 1×10^5^ of B16F0 cells in 300 μl PBS were inoculated subcutaneously (s.c) in the right flank.

On days 12, 13, 15 and 17 of tumor growth mice were treated with 250 μg anti-CTLA4 (Clone 9D9, Bio X Cell #BP0164, USA) injected intraperitoneally (i.p.). Tumor sizes were measured with a caliper three times a week. Tumor volume was calculated using the formula V=a*b*1/2b, where a and b are larger and smaller lateral dimensions of the tumor. On days 19 and 20 mice were euthanized with Isoflurane, inguinal lymph nodes from the tumor flank and from the opposite flank were scissored out. Lymph nodes were dissected and treated with the Liberase/DNAse solution as described above.

### FACS sorting

Cells from both popliteal LNs were combined and stained with the following antibodies anti-CD3-APC (clone 17A2), anti-CD19-PE/Cy7 (Clone 1D3), anti-CD4-V450 (Clone RM4-5), anti-CD8-APC/Cy7 (Clone 53-6.7) all from BD Biosciences. Staining was done in PBS-BSA: PBS supplemented with 1% bovine serum albumin (BSA, Sigma-Aldrich) on ice for 1 h. After 1h stinging on ice cells were washed, resuspended in PBS-BSA buffer at 5-10×10^6^ cells/ml, analyzed and sorted with FACSAria III cell sorter (BD Biosciences) in 2 ml RPMI cell culture medium (Gibco, USA) supplemented with 10% FBS. 10^6^ Th and CD8 and 1-5×10^5^ Treg cells were sorted for each mouse. Sorted cells were washed twice with PBS, lysed with 300 μl RLT buffer (Qiagen, Hilden, Germany) and freeze and stored in this buffer at −80°C before RNA extraction.

Cells from inguinal LNs of tumor bearing mice were stained with by the same protocol and with same antibodies as for popliteal LNs but with addition of anti-CD45-PerCP/Cy5.5 (clone 30-F11), anti-CD25-PE (clone PC61), CD69-BV510 (clone H1.2F3) all from BD Biosciences. Unlike with popliteal LNs only one LN was used for each probe and these LNs were not as enlarged as popliteal LNs after vaccination. As a result there were 6-20×10^4^ CD8+ and Th cells in each probe and cells were sorted directly in 300 ul of RLT buffer with the limit of 7.5×10^4^ cells per tube. Right after sorting tubes were vortexed left at RT for at least 10 min to complete lysis. After lysis cells were stored at −80°C before RNA extraction. Single tubes with 6-7.5×10^4^ cells were used for further TCR or RNA sequencing.

All the experiments conducted with animals were carried out in accordance with the National Institutes of Health guide for the care and use of Laboratory animals (NIH Publications No. 8023, revised 1978). The experimental protocol was approved by the Ethical Committee of the Nizhny Novgorod State Medical Academy (EC #11, granted March 12, 2015).

### TCR sequencing

Cell samples in the RLT buffer were thawed and RNA was extracted using RNeasy Mini Kit (Qiagen). T-cell receptor beta chain (TCRβ) cDNA libraries were prepared using the Mouse TCR RNA kit (MiLaboratories Inc) according to the manufacturer’s protocol. Briefly, cDNA libraries were synthesized using 5’RACE (Rapid Amplification of cDNA Ends) with unique molecular identifiers (UMI) incorporated within template switch oligo, TCR-specific primers and SMARTScribe Reverse Transcriptase (Takara Inc). After purification with Agencourt AMPure XP magnetic beads (Beckman Coulter, Beverly, MA) cDNA samples were amplified with 19 cycles of PCR with Q5® Hot Start High-Fidelity DNA Polymerase (New England Biolabs) and supplied primes. PCR product purification was purified with AMPure XP magnetic beads and used for second PCR with supplied primers and IDT® for Illumina® DNA/RNA UD Indexes (Illumina). After 10-11 PCR cycles, libraries were pooled in a single sample, purified with AMPure XP magnetic beads and checked for the appropriate length (600-700nt) using TapeStation capillary electrophoresis (Agilent Technologies). Libraries were sequenced with 150+150 nt sequencing kits on MiSeq or NextSeq platforms (Illumina) with 10^6^ reads per sample in peptide vaccination experiments or 10-30 reads per sorted T-cell in CTLA-4 experiment.

### RNA sequencing

For transcriptome RNA sequencing cells were sorted in the RLT buffer and RNA was isolated as described above. cDNA were prepared with SMART-Seq® v4 Ultra® Low Input RNA Kit for Sequencing (Tacara Bio, Japan) according to manufacturer’s instructions. Further library preparation was done with Nextera XT DNA Library Preparation Kit (Illumina). Sequencing was performed on NextSeq platform (Illumina) using 150+150 nt sequencing kits with 10^7^ reads per sample.

### Data Analysis

Repertoires were extracted from RNA-based TCRβ sequencing or from RNA-Seq data with MiXCR software.

Pooling data and TCRnet are described in VDJtools documentation (https://vdjtools-doc.readthedocs.io/en/master/) as *JoinSamples* and *CalcDegreeStats* functions. Getting the top of clonotypes was done by python script.

For TCRnet we used control TCRβ CDR3 repertoires that were randomly generated by OLGA pipeline `olga-generate_sequences --mouseTRB` (https://github.com/statbiophys/OLGA). Pipeline for cluster construction was used as described (https://github.com/antigenomics/tcr-annotation-methodology/blob/master/tutorial.Rmd).

All statistics was done in R. We used T-tests from `ggpubr` library in R and One-way ANOVA with comparison for each group within the group-mean based on Wilcoxon test (or comparison for each group versus control group). P-Value is shown as: * p < 0.05, ** p < 0.01.

All the graphs were created in R using the ggplot2 library. Boxplots correspond to the 25 to 75 interquartile range with the median value. Extreme points outside of the minimum/maximum values are also shown. `ggplot2` boxplots shows 75 percentile as upper box, 50th percentile (median) as line in the box, and 25th percentile as bottom of the box. Upper and lower whiskers show the largest and the smallest value of the data. Outside value is 1.5 times >/< the interquartile range beyond either end of the box.

### Data availability

All data is available in Figshare: https://figshare.com/account/articles/24637356. Dataset of identified B16 neoantigen-specific TCRs is available in VDJdb database https://vdjdb.cdr3.net/search repository, https://github.com/antigenomics/vdjdb-db/issues/397.

## Results

### Identification of TCRβ CDR3 motifs recognizing specific B16 melanoma peptides

To identify public TCRβ CDR3 motifs reproducibly involved in response to particular B16 melanoma neoantigenic peptides in C57BL/6 mice, we used vaccination of mice groups, followed by extraction and sorting of T cell subsets from the draining lymph node, TCRβ CDR3 repertoire profiling, and computational extraction of TCRβ CDR3 motifs reproducibly enriched in response to each particular peptide compared to any other peptide. Since the C57BL/6 strain of mice expresses only a single MHCII molecule, I-Ab, this allowed us to assign the identified CD4+ TCRβ CDR3 motifs to this particular MHC-II variant.

14 immunogenic B16F10 peptides carrying somatic mutations that were shown to have therapeutic activity and/or induce prominent *in vitro* reactivity (**Table 1**) were selected from the keystone studies of Ugur Sahin and colleagues (6) (7). In these works, non-synonymous mutations of the B16F10 tumor were mapped, and resulting mutated peptides were screened for immunogenicity using several approaches. Most of the tested long peptides, including those that contained MHCI-presentable short peptides within (12), previously elicited helper CD4+ but not CD8+ T cell response, both in peptide and mRNA-based vaccination assays. Conceptually, these works have shown importance of neoantigen-specific CD4+ T cell response in tumor control and efficiency of immunotherapeutic and vaccination interventions (5,7,13,14).

**Table 1.**
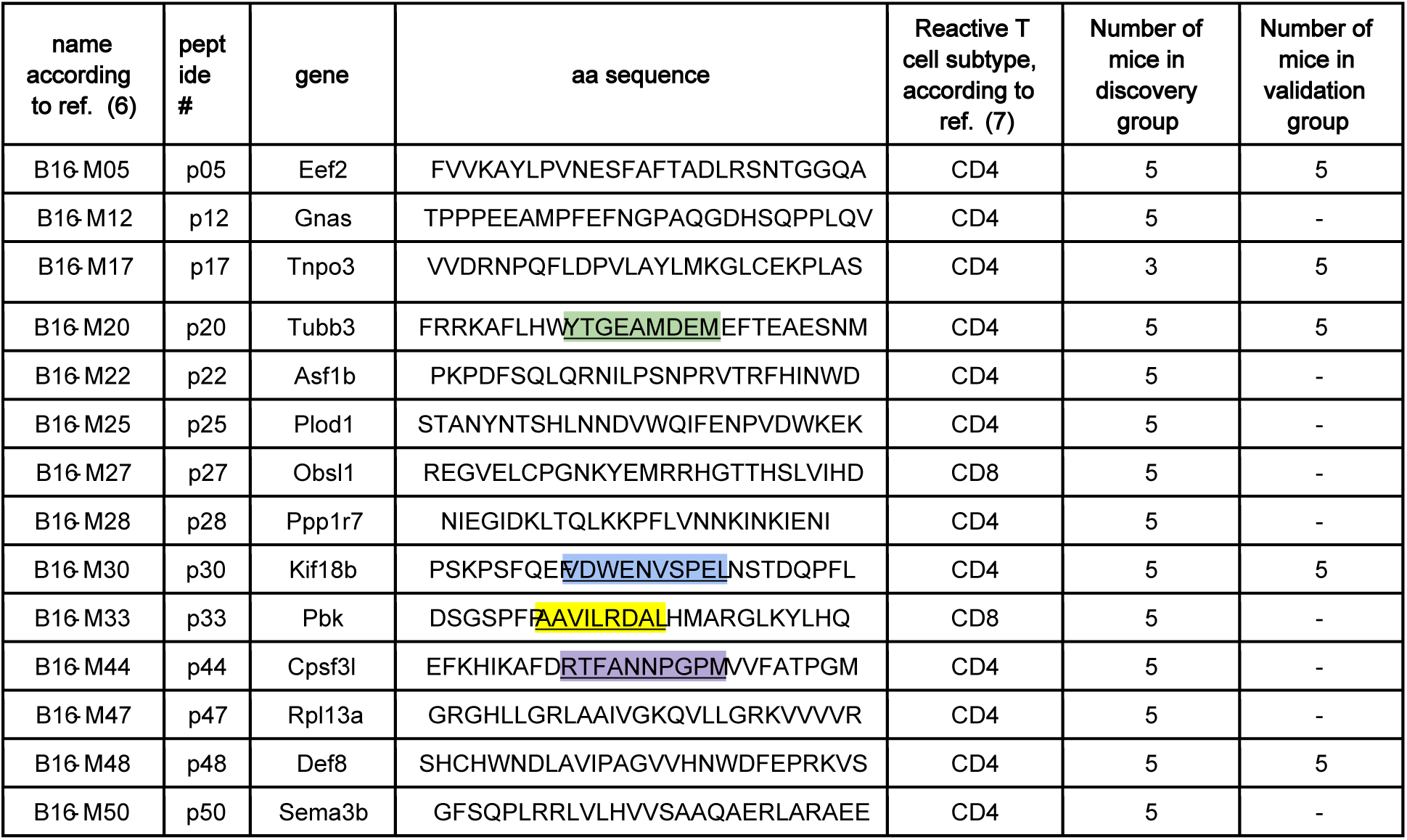
B16F0 peptides and mice groups used in the study. B16F0 peptides and mice groups used in the study. Short peptides contained within the long peptides that can be presented in H2-K^b^ or H2-D^b^ MHCI context of C57Bl/6 mice are underlined.

Here we tested the ability of those peptides to elicit clonal expansions within CD4+, Treg, and CD8+ T cells that would share common TCRβ CDR3 motifs across vaccinated animals.

For these experiments, we used C57Bl/6-FoxP3^EGFP^ mouse strain. In the first experiment, “discovery” groups of 5 mice each underwent footpad vaccination with 20 mg of one of the 14 peptides in each heel, in incomplete Freund’s adjuvant (IFA). After 2-3 weeks popliteal lymph nodes were taken, and CD8+, CD4+ Th, and Treg subsets were sorted by FACS (**Supplementary Fig. S1a, Supplementary Table 1**). TCRβ cDNA libraries were prepared using the 5’RACE approach as described(15). TCRβ CDR3 repertoires were extracted using MiXCR software, MiLaboratories Inc (https://milaboratories.com/). Downstream analysis was performed using VDJtools(16) and R.

To identify antigen-specific TCR clusters, we pooled the top-1000 most frequent amino acid TCRβ CDR3 clonotype sequences from each mouse for each group of 5 mice vaccinated with the same peptide. We next used TCRnet feature of VDJtools (17) to identify amino acid CDR3 variants with more neighbors (Hamming distance = 1, same TRBV segment) than expected by chance. For this, control TCRβ CDR3 repertoires were randomly generated by OLGA pipeline (`olga-generate_sequences`, https://github.com/statbiophys/OLGA). Resulting TCR clusters were built from the core CDR3β variants and their neighbors (**Fig. 2, Supplementary Table 1**).

**Figure 2.**
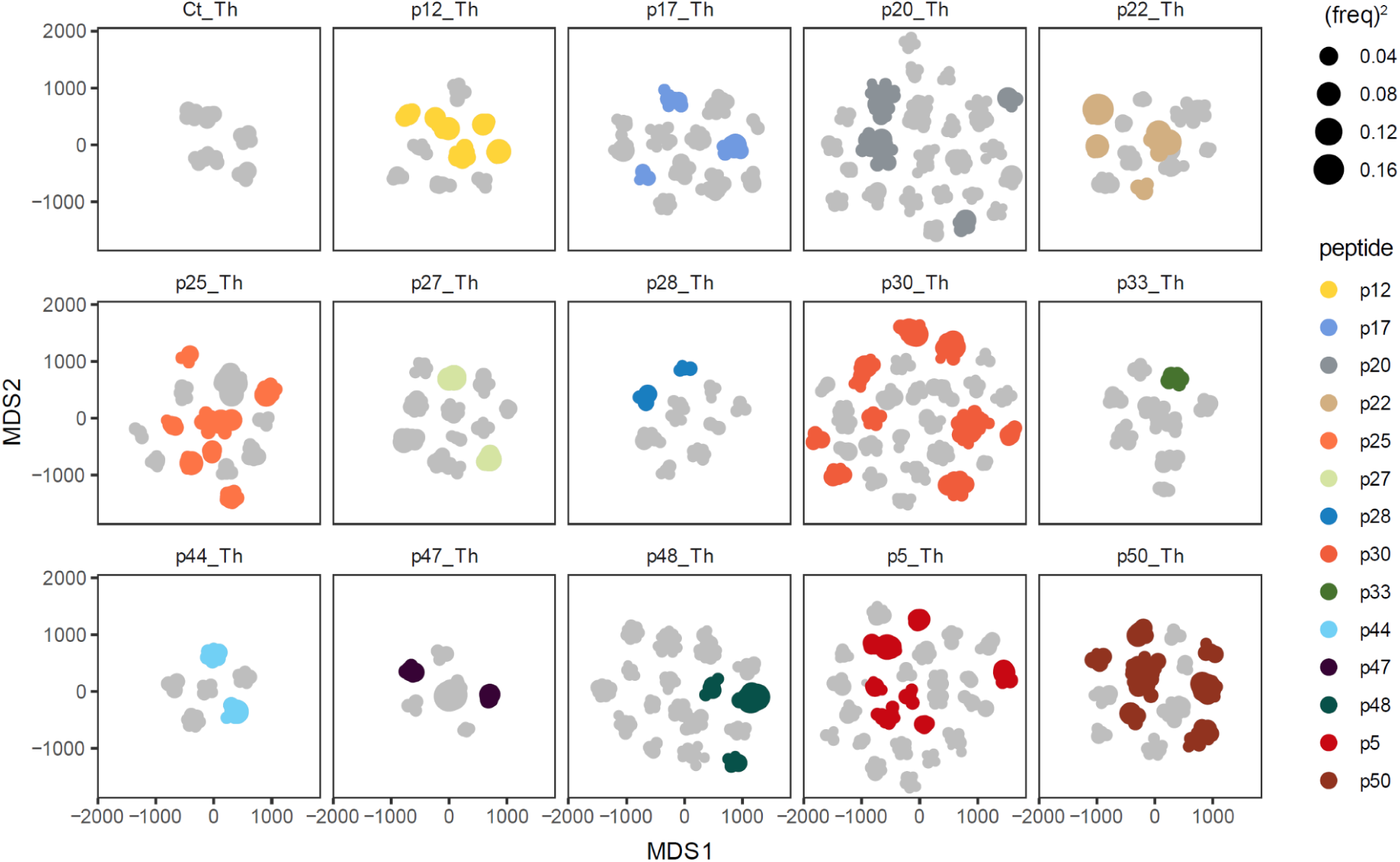
Identified TCRβ CDR3 clusters specific for B16 antigens. Clusters were identified using TCRnet from the most common clonotypes of the group of mice vaccinated with a single peptide. Each cluster is shown as a node. The specific TCRβ CDR3 clusters selected by edgeR are colored, while the gray represents the clusters that did not pass edgeR selection.

Next, we used edgeR package in R to identify TCR clusters that were selectively enriched in mice vaccinated with the same peptide, comparing them pairwise with each group of mice vaccinated with a different peptide (https://bioconductor.org/packages/release/bioc/html/edgeR.html). We used glm approach: quasi-likelihood F tests for pairwise comparison in edgeR. Cumulative UMI counts of each TCR cluster were analyzed. TCR clusters that passed all 13 peptide-peptide comparisons with the p-value < 0.05 were selected. Additionally, only TCR clusters represented in at least 2 mice of the group were selected. These clusters are represented on **Fig. 2** as colored clusters.

Abundance of selected TCR clusters among mice that got different peptide vaccinations is shown on **Fig. 3**. On this heatmap we show summary frequency of clonotypes which entered the cluster with one amino acid mismatch allowed within CDR3aa, TRBV segments fixed.

**Figure 3.**
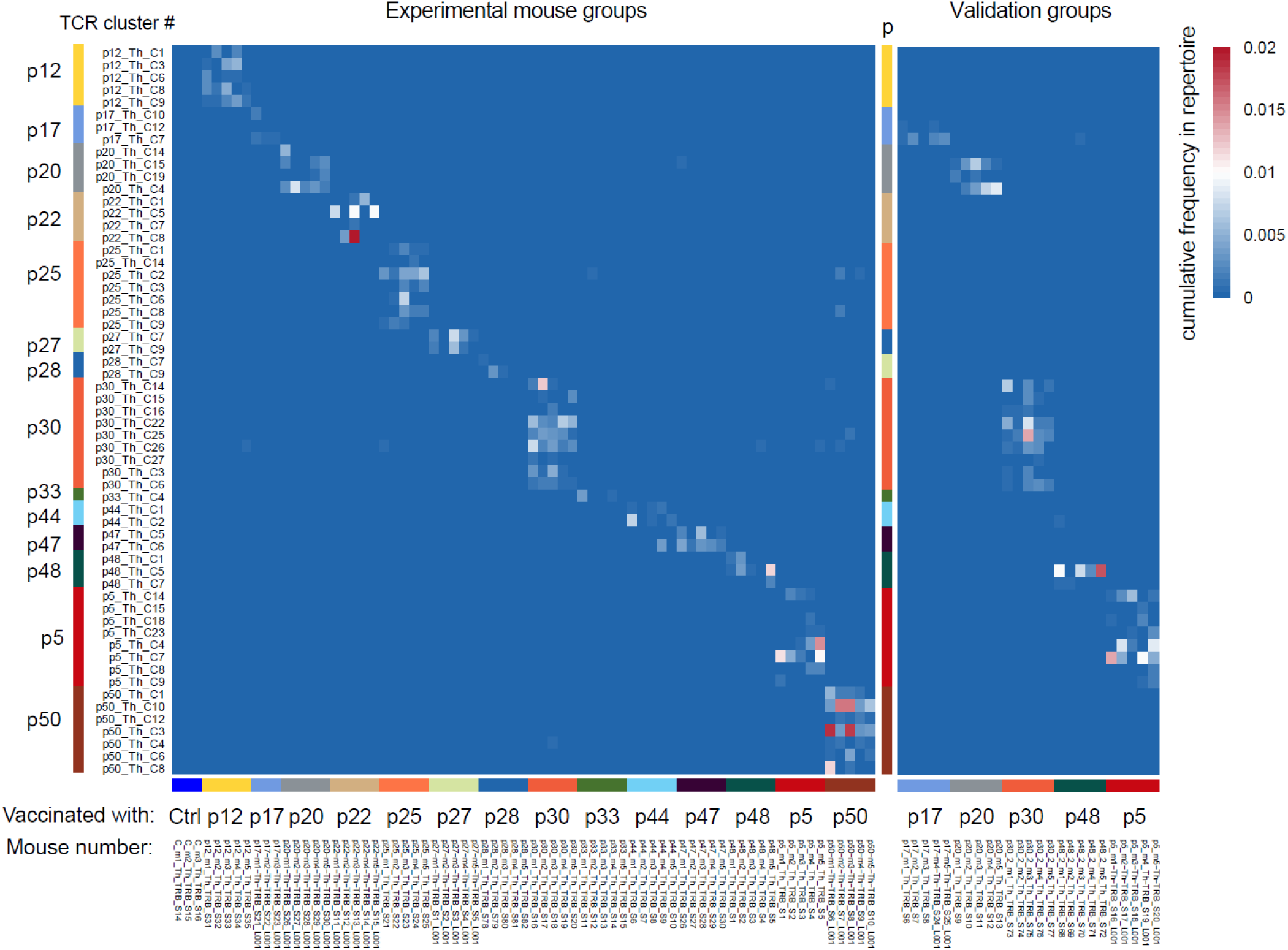
Distribution of identified B16F10-specific Th TCRβ CDR3 clusters. Cumulative frequencies of all TCRβ CDR3s from each TCR cluster (rows) in each mice (columns) are shown for CD4+ TCRβ CDR3 repertoires from discovery and validation mice groups.

**Figure. 4.**
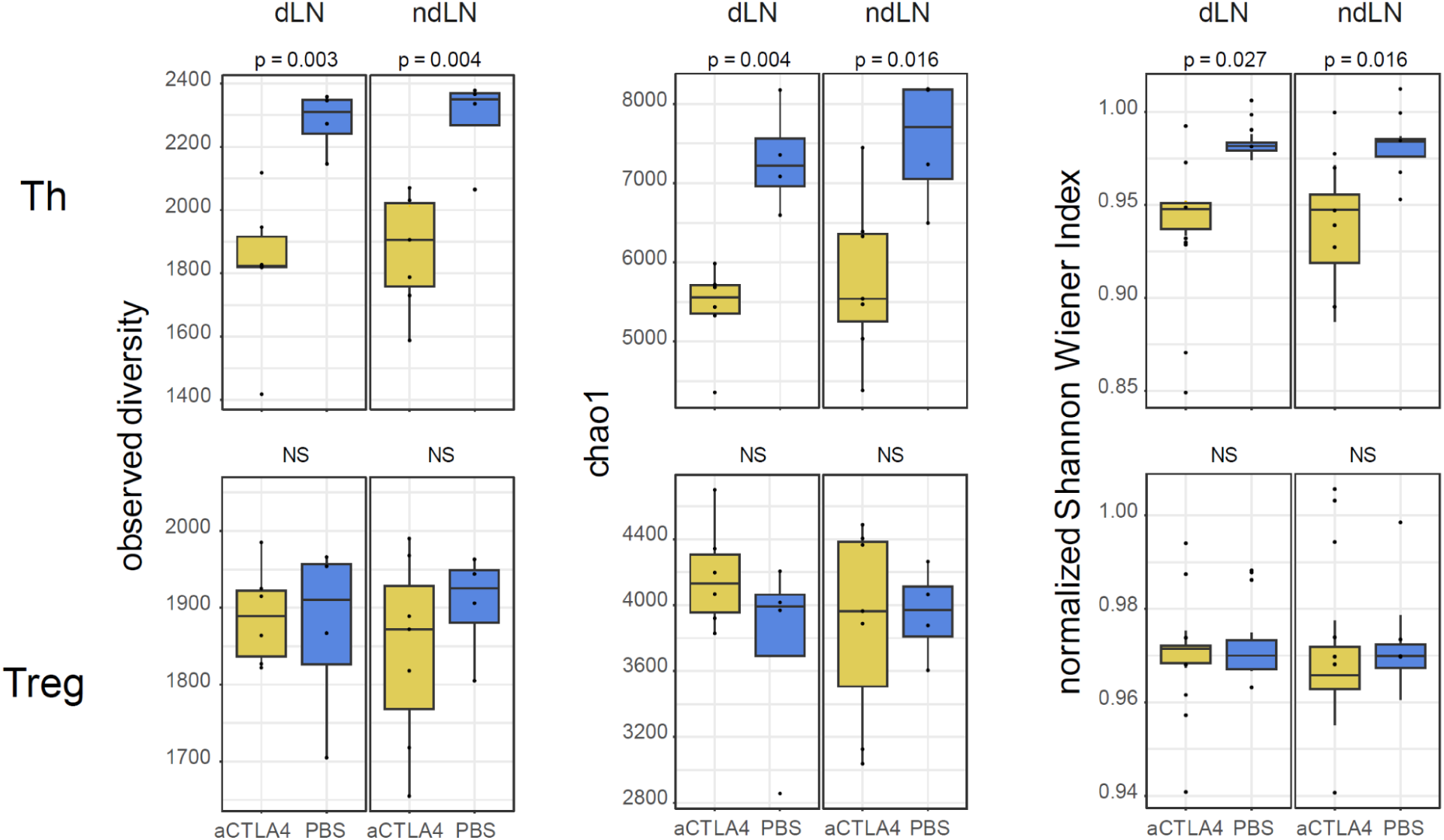
Th and Treg TCRβ CDR3 repertoire diversity in dLNs and ndLNs of anti-CTLA4 treated and control mice. Observed diversity (number of clonotypes), Chao 1 (lower diversity estimate) and normalized Shannon-Wiener indexes (inverse of clonality, reflects evenness of clonotype size distribution) metrics are shown. For normalization, each cloneset was preliminary down-sampled to 3000 randomly chosen CDR3-containing sequencing reads. P-values based on parametric t-test for CTLA4 treatment group versus PBS control are shown. Boxplots correspond to the 25 to 75 interquartile range with the median value. Extreme points outside of the minimum/maximum values are also shown.

To confirm the validity of this approach for selection of peptide-specific TCR clusters, we analyzed abundance of the same clusters on additional validation groups of mice (**Table 1**), vaccinated with peptides p5, p17, p20, p30, and p48. This analysis confirmed enrichment of relevant TCR clusters exclusively in mice vaccinated with corresponding peptides (**Fig. 3**).

For each CD4+ T cell TCR cluster, we now had information on included CDR3β sequences, cognate peptide, and MHCII, since C57BL/6 mice carry only I-A^b^/I-E^b^ MHC II. Altogether, we extracted 59 TCRβ CDR3 clusters that include 485 TCRβ CDR3 amino acid and 1556 TCRβ CDR3 nucleotide sequence variants as summarized in **Supplementary Table 3**. See also **Supplementary Fig. S2** for CDR3 logo images.

Identified B16-specific C57BL/6 TCRβ CDR3 sequence variants were uploaded to the VDJdb database (https://vdjdb.cdr3.net/) and can now be used in vaccine and immunotherapy studies on C57BL/6-B16 tumor models.

Of note, the same analysis pipeline extracted no reproducibly observed TCRβ CDR3 clusters in Treg and CD8+ T cell subsets. For Tregs, this may be interpreted as absence of prominent antigen-specific priming in our adjuvant peptides vaccine setting. Predominantly CD4 but not CD8 response to the long peptides vaccination was previously shown and expected (6,7).

To investigate potential plasticity between Th and Treg subsets for the peptide-responding T cell clones, we mapped identified B16-specific Th TCRβ CDR3 clusters on Th and Treg repertoires, allowing additionally for 1 aa mismatch (**Supplementary Fig. S3**). In general, Th clusters were represented poorly and irregularly in Treg repertoires. At the same time, some clusters showed more regular overlap between CD4+ and Treg subsets of corresponding mice groups, suggesting non-zero plasticity/convergence/asymmetric naive T cells priming between Th and Treg subsets (shown by rectangles on **Supplementary Fig. S3b**).

### Tracking B16-specific T cell clones in response to blocking anti-CTLA4 immunotherapy

Next, we used the obtained database of B16 peptides-specific TCRβ CDR3 motifs to track the behavior of B16-specific T cell clones in response to blocking anti-CTLA4 immunotherapy on the orthotopic model of B16F0 melanoma (Teicher BA. *Tumor Models in Cancer Research.* Boston, USA: Springer; 2010. p. 682.). We used noncytotoxic, blocking only, mild action anti-CTLA4 antibody clone (9D9, IgG2b, BioXcell #BP0164 (18)). It has modest therapeutic efficiency that is thought to be compromised by some protumor activities. The latter include expansion of Tregs in the periphery and in lymph nodes and promotion of intratumoral and peripheral PD-1^high^ follicular T cells (19,20). In order to dissect tumor-specific responses, we analyzed sorted Th and Tregs from tumor draining (dLN) and non-draining (ndLN) inguinal lymph nodes.

We used 8-11 weeks old male hemizygotic C57BL/6-FoxP3^EGFP^ mice model, inoculated subcutaneously with B16F0 cells. Mice were treated with anti-CTLA4 antibodies (n=7) or PBS (n=4) at days 12, 13, 16 and 17. Slight but not statistically significant suppression of tumor growth was observed after the first two treatment doses, and this suppression was completely abolished at later stages (**Supplementary Fig. S4**).

Tumor draining lymph nodes (dLNs) and non-draining lymph nodes (ndLNs) were isolated on days 19 and 20. LNs were dissociated into single-cell suspensions and stained for FACS sorting. CD4, CD8 and Treg cells were gated (**Supplementary Fig. S1b**) and sorted directly in the RLT Lysis buffer (Qiagen) to preserve RNA for further extraction.

RNA-Seq was performed using SMART-Seq v4 Ultra Low Input RNA Kit, Takara Bio (transcriptome analysis to be published elsewhere). TCRβ CDR3 repertoires were extracted from RNA-Seq data using appropriate preset parameters of MiXCR software (21,22). Altogether, we obtained TCRβ CDR3 repertoires for Th and Treg cells from 8 LNs samples of 4 control mice (dLN and ndLN for each) and 13 LNs of 7 anti-CTLA 4-treated mice (no LN1 in mice #3 due to technical sample loss). Each Th repertoire contained 4847±606 (median±s.d.) TCRβ CDR3 reads with 2885±404 unique clonotypes. Each Treg repertoire contained 4520±882 TCRβ CDR3 reads with 2555+332 clonotypes.

#### Repertoire diversity and clonality

We estimated TCRβ CDR3 repertoire diversity and clonality using previously evaluated metrics, including observed diversity (number of distinct clonotypes), normalized Shannon Wiener index (repertoire evenness and the extent of clonal expansion) and Chao 1 (estimated lower bound of total diversity based on relative representation of small clonotypes) (15). We found decrease of all diversity metrics in anti-CTLA4 treatment group in the case of Th but not of Treg cells which indicates therapy-induced increase in the Th clonality (relative representation of large clonal expansions).

#### Tracking B16 peptide-specific T cell clonotypes

Next, we analyzed relative abundance of the identified B16-peptide specific TCRβ CDR3 clonotypes in dLNs and ndLNs of CTLA-4 treated and control mice. We matched peptide-specific amino acid CDR3 clоnotypes with Th and Treg TCR repertoires, allowing for single amino acid mismatch, TRBV segment fixed. Overall, corresponding B16-specific clonotypes were represented at comparable levels in both LNs of both treated and non-treated mice (**Fig. 5a**). Remarkably, and quite unexpectedly, B16-peptide specific TCRβ CDR3 clonotypes, initially identified for Th (**Figs. 2,3**) but absent in Treg repertoires of vaccinated mice (**Supplementary Fig. S3b**), were well represented among Tregs obtained from LNs of tumor-bearing mice (**Fig. 5b**).

**Figure 5.**
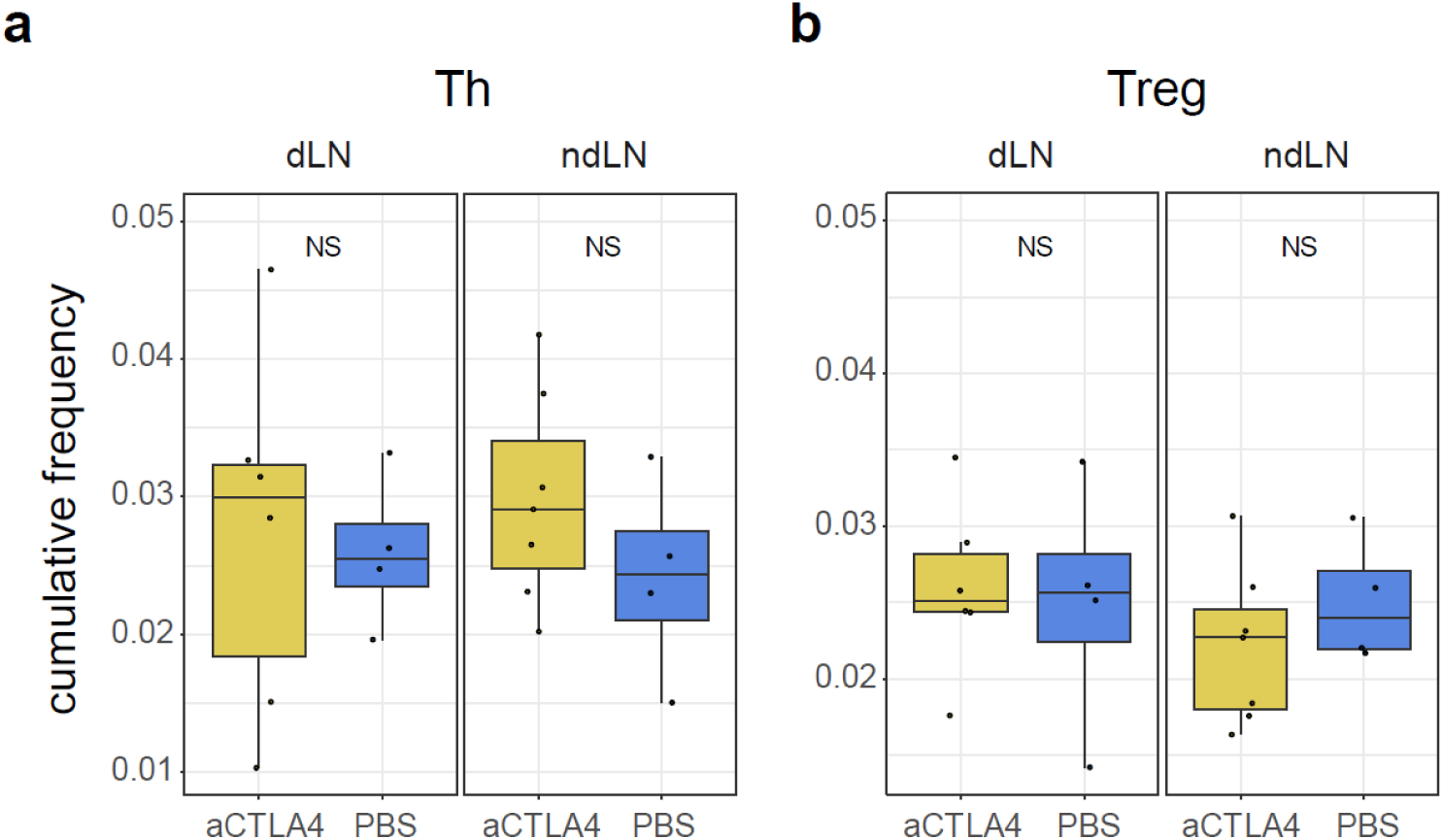
Relative abundance of B16-peptide specific TCRβ CDR3 clonotypes in dLNs and ndLNs repertoires. Plots show cumulative frequency of clonotypes that match B16-peptide specific TCRβ CDR3 clonotypes with allowed single amino acid mismatch, TRBV segment fixed, within Th and Treg TCRβ CDR3 repertoires of dLNs and ndLNs of mice treated with aCTLA4 or PBS. Boxplots correspond to the 25 to 75 interquartile range with the median value. Extreme points outside of the minimum/maximum values are also shown.

We next examined the specific response for each peptide. At the level of peptide-specific groups of TCR clusters, LN repertoires of treated and non-treated mice also looked comparable (**Fig. 6a,b**, 1mm RNA-Seq, boxplots separate for each peptide). Response to particular peptides, including p5, p20, p25, p30, and p50 was reproducibly more prominent across LNs and mouse groups, indicating their higher immunogenicity. Again, B16-specific TCR clusters were well represented in Treg repertoires, with the patterns similar to Th subset (**Fig. 6c,d)**.

**Figure 6.**
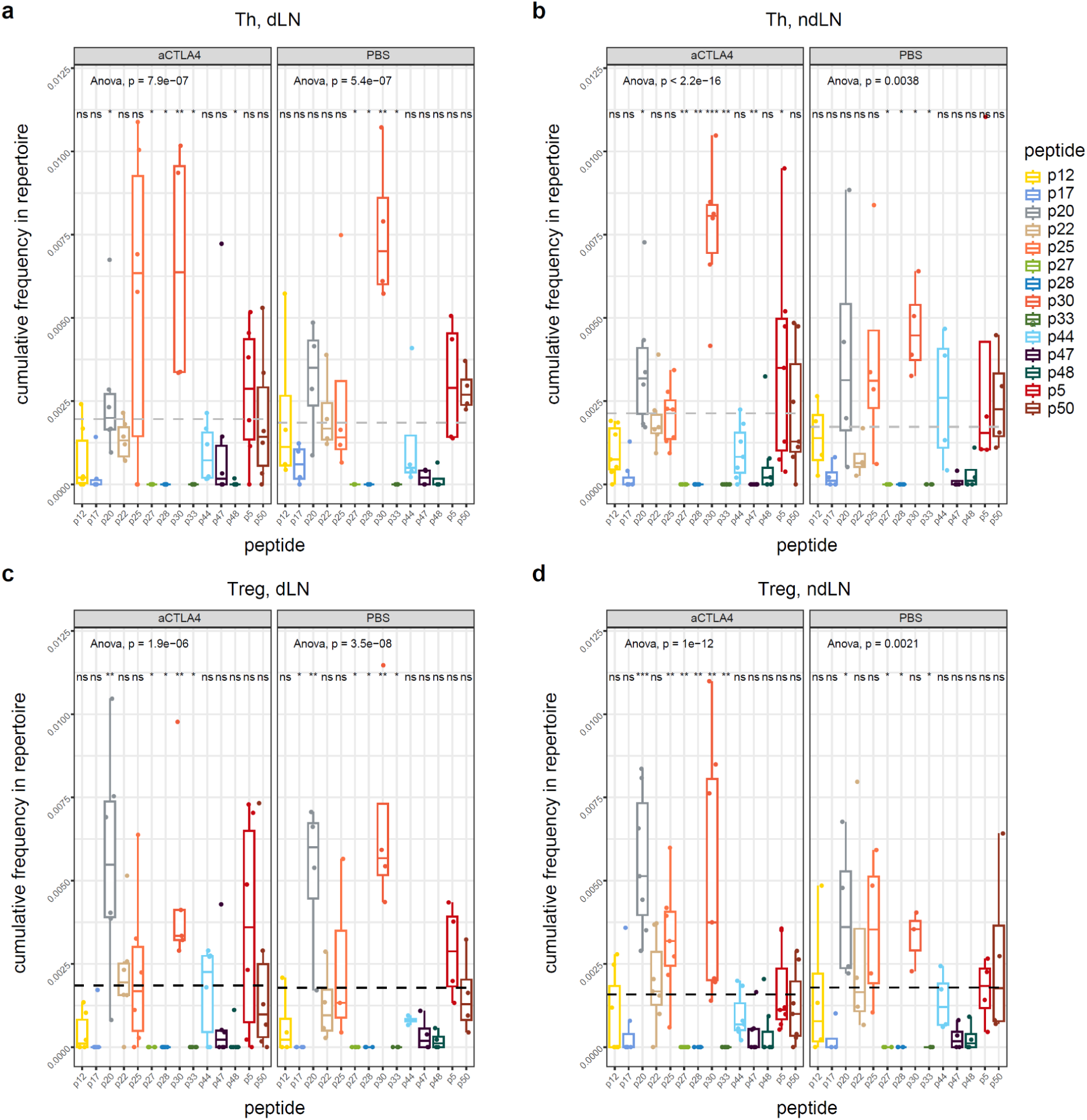
Cumulative frequencies of B16 peptide-specific clonotypes. Each point corresponds to abundance of TCRs specific to one peptide in the repertoire of one mouse. Clonotypes were matched on the basis of identical TRBV gene segments and TCRβ CDR3 aminoacid sequences with 1 mismatch allowed. Data shown for Th (a,b) and Treg (c,d); for dLNs (a,c) and ndLNs (b,d). One-way ANOVA p value is shown for each panel. Comparisons for each peptide specificity within the group-mean based on Wilcoxon test are shown as * p < 0.05, ** p < 0.01. Boxplots correspond to the 25 to 75 interquartile range with the median value. Extreme points outside of the minimum/maximum values are also shown.

Similar picture was observed at the level of individual TCR clusters, but here cluster p30_Th_C26 was overrepresented in anti-CTLA4 treated animals (**Fig. 7**).

**Figure 7.**
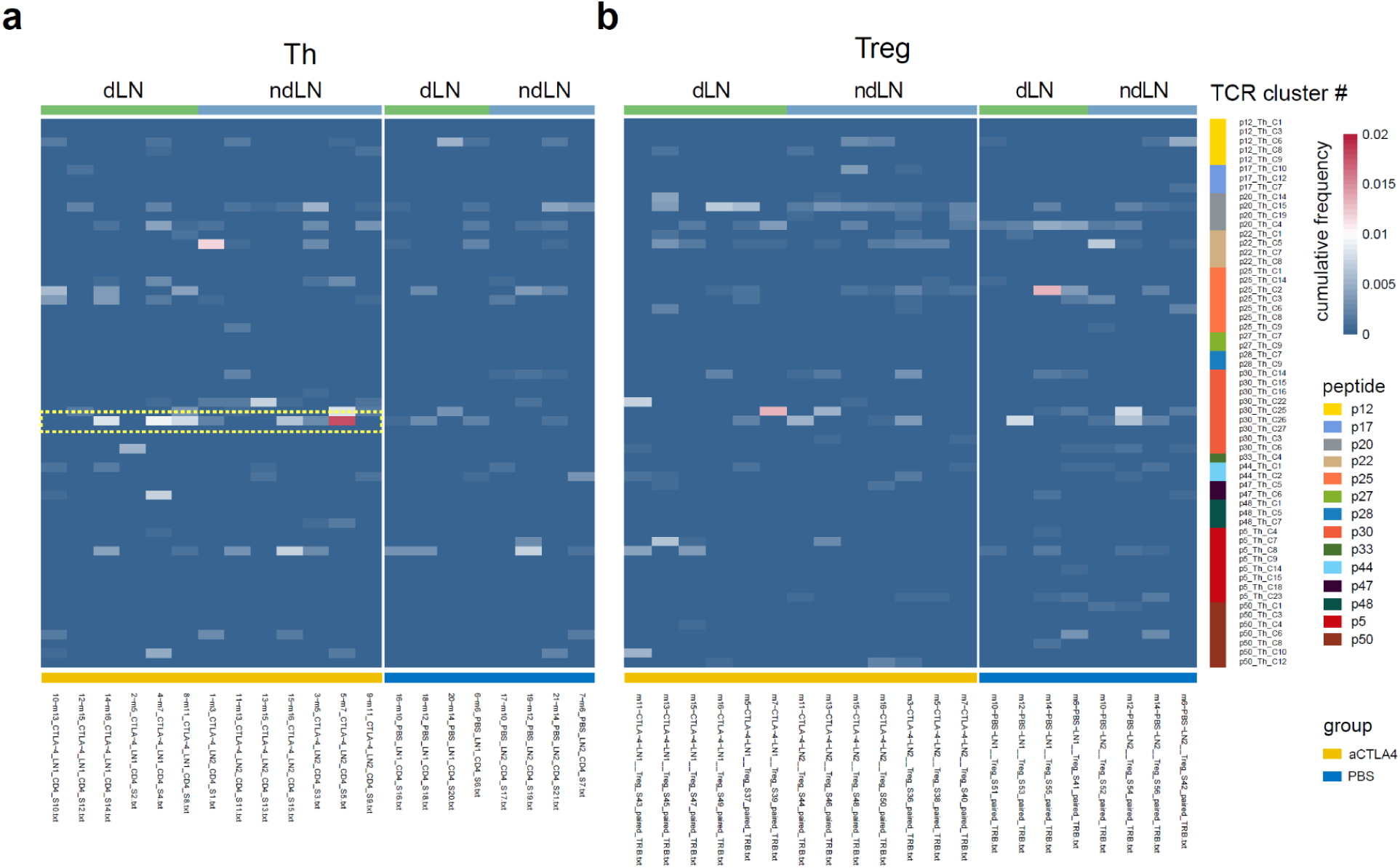
Frequencies of B16 peptide-specific TCR clusters. Heatmap shows abundance of each B16 peptide-specific TCRβ CDR3 cluster in Th (a) or Treg (b) repertoires for dLNs and ndLNs. Each mouse repertoire is shown in a separate column. Clonotypes were matched on the basis of identical TRBV gene segments and TCRβ CDR3 aminoacid sequences with 1 mismatch allowed.

#### Th to Treg plasticity

To assess the possible clonal conversion of Th to Treg, we analyzed the overlap of the top-1000 clonotypes from each TCRβ repertoire at the CDR3 nucleotide level, for the paired Th-Treg samples obtained from the same mice. We used F2 repertoire similarity metrics of VDJtools software, which employs a clonotype-wise sum of geometric mean frequencies that takes into account the relative size of shared clonotypes. The closer two samples are, the higher the F2 metric, which reflects the overall frequency of shared clonotypes(15,23). We also used the D metric of VDJtools which measures the number of shared clonotypes between the two samples. The relative repertoire overlap between Th and Treg nucleotide CDR3 repertoires was notably higher in anti-CTLA4 treated mice (**Fig. 8**). This may indicate that in the context of aCTLA4 suppressed Treg function, Th clones acquire Treg phenotype as a compensatory mechanism, most probably temporarily(23).

**Figure 8.**
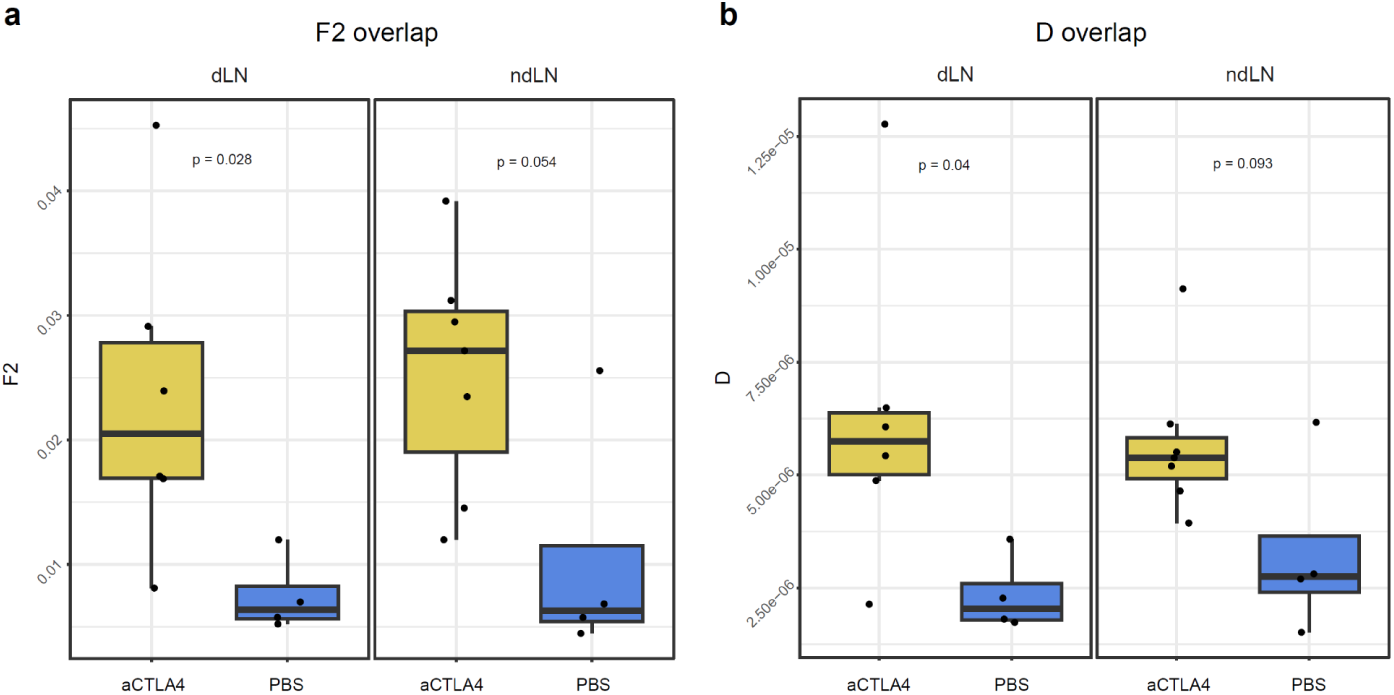
Th to Treg plasticity. Nucleotide TCRβ CDR3 repertoires overlap measured by F2 (a) and D (b) VDJtools metrics are shown. Overlaps measured for top-3000 clonotypes from each repertoire, for paired Th-Treg repertoires obtained from the same mice, same LN. Boxplots correspond to the 25 to 75 interquartile range with the median value. Extreme points outside of the minimum/maximum values are also shown.

Importantly, analysis of these Th-Treg overlapped nucleotide clonotypes identified no TCRβ CDR3 variants specific to B16 melanoma peptides. This indicates that prominent representation of B16 melanoma-specific TCRβ CDR3 variants in Treg repertoires of tumor-bearing mice shown above (**Figs. 5,6,7**) does not result from Th to Treg plasticity that is mainly induced by aCTLA4 treatment (**Fig. 8**), but is rather explained by independent priming of convergent Th and Treg clones(11) specific to the same cancer neoantigens.

In other words, we observe that Th and Treg clones share the same tumor neoantigens specificity (highly similar or identical amino acid CDR3 variants), yet independent clonal origin (distinct nucleotide CDR3 sequences). It is also important to underline that B16 melanoma-specific TCRβ CDR3 variants were nearly absent in Treg repertoires of peptide-vaccinated mice (**Supplementary Fig. 3**) yet appear in LNs of tumor bearing mice (**Figs. 5,6,7**). Altogether, our observations suggest tumor influence on dendritic cells that promotes priming and expansion of tumor-specific Treg clones.

#### Analysis of TCR clusters derived from tumor-bearing mice

Finally, we analyzed TCRβ CDR3 clusters derived from the Th and Treg repertoires of LNs obtained from tumor-bearing mice. For Treg and CD4 repertoires separately, we pooled the top-500 most frequent amino acid CDR3β clonotype sequences from LN repertoires of 4 mice treated with CTLA4 and 4 mice treated with PBS (16 repertoires, 8,000 clonotypes for Th and 16 repertoires, 8,000 clonotypes for Treg subsets). Using TCRnet and control repertoires that were randomly generated by Olga, we built clusters from the core TCRβ CDR3 variants and their neighbors (Hamming distance = 1, same TRBV segment) among all repertoires.

Notably, the newly identified Th TCR clusters were more prominently represented in Th repertoires of anti-CTLA4 treated mice compared to PBS-treated animals (**Fig. 9**). Obtained Th clusters included TCRβ CDR3 variants specific to melanoma peptides (p20, p30, and p50), that were better represented in Th clusters of anti-CTLA4 treated mice compared to PBS controls (**Fig. 9**). In contrast, melanoma-specific TCRβ CDR3 variants were better represented in Treg-derived clusters of PBS-treated mice (**Supplementary Fig. S5**).

**Figure 9.**
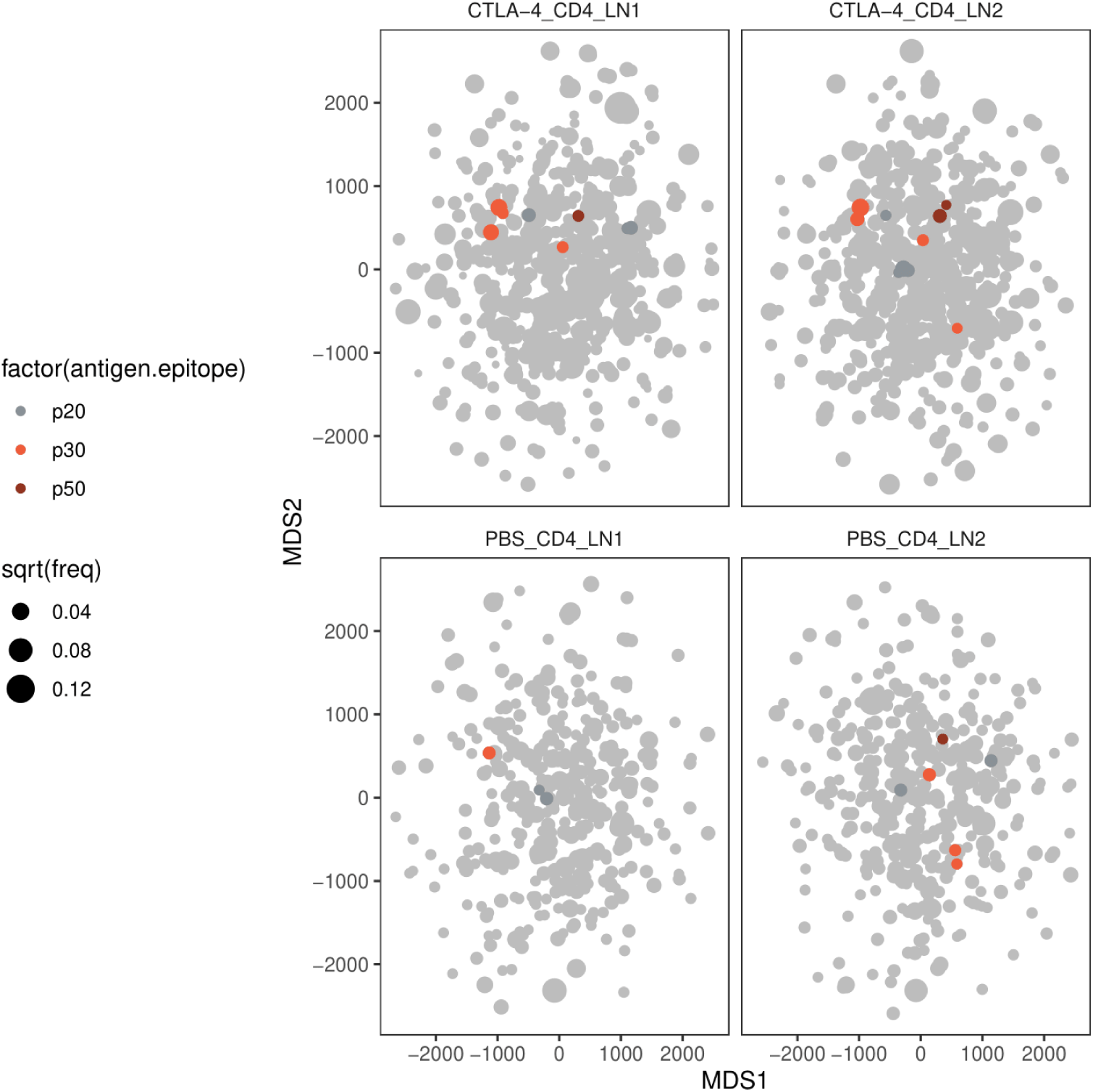
Th TCRβ CDR3 clusters derived from tumor-bearing mice. Clusters were identified using TCRnet from the integrated top-500 clonotypes from Th TCRβ CDR3 repertoires from dLNs and ndLNs of PBS and anti-CTLA4 treated mice. Each cluster is shown as a node. Clusters that match identified melanoma peptide-specific TCRβ CDR3 variants are colored.

## Discussion

Although tumor-specific T cells usually represent a relatively minor proportion of all tumor infiltrating lymphocytes (24), their presence and functional activity is well established as a critical feature of anti-tumor response and efficiency of immunotherapeutic and vaccination approaches (25). Among those, helper T cells play a plethora of organizational roles as antigen-specific managers of immune response (5,26,27).

Ability to track for the fate, abundances, functionality, and subset attribution of T cells clones specific for characteristic tumor neoantigens should greatly facilitate vaccine and immunotherapy development on animal models. Convergent shared antigen-specific TCR motifs(11) provide an excellent opportunity to facilitate such studies.

Here we identified convergent clusters of TCRβ CDR3 variants specific to 14 previously described B16 melanoma neoantigenic peptides in I-Ab pMHCII context of C57BL/6 mice. These data are available in VDJdb (https://vdjdb.cdr3.net/) and can be employed for informed tracking of tumor-specific T cell cones in cancer research and development studies.

We next employed this database to track B16-specific T cell clones on the orthotopic melanoma model, to investigate the action mechanics of blocking anti-CTLA4 antibody.

In humans, anti-CTLA4 therapy with ipilimumab acts through blocking of CTLA and depletion of Tregs, the cell subset with the highest CTLA4 expression (28–30). The primary mechanism for efficacy of anti-CTLA4 is FcR-mediated Treg depletion in the tumor microenvironment. Elimination of Tregs via ADCC and complement activation may result in activation of antitumor immune response but is associated with systemic autoimmunity (31–33).

Despite numerous pre-clinical mouse studies showing that CTLA4-blocking antibodies with appropriate Fc domains could mechanistically deplete regulatory T cells (Tregs) in regressing tumors, data remains scarce in humans associating this with clinical response (34). In mice there are two commonly used anti-CTLA4 clones, 9H10 and 9D9 (19,35). If the first one depletes Treg, here we have used the 9D9 clone that possesses only blocking activity and has lower therapeutic efficiency compared to 9H10 (19,36). We have observed only minor tumor suppression and only at the early stage of the therapy (**Supplementary Fig. S5**). The reasons for low efficiency of CTLA4-blocking remains understudied. Among possible reasons, glucose deprivation in tumor environment, stimulation of peripheral Treg and activation of suppressive phenotype of follicular T cells are considered (19,20,37).

Here, on the orthotopic B16 melanoma model we observed that CTLA4-blocking therapy results in prominent increase of clonality in Th but not Treg subset in both dLNs and ndLNs, reflecting unblocked clonal expansion of helper T cell clones in the absence of mainly Treg-derived CTLA4-mediated pressure (38–40).

Identified TCRβ CDR3 variants specific to B16 melanoma neoantigens were comparably represented in both Th and Treg repertoires, both in anti-CTLA4 treated and control mice. This observation was quite remarkable, considering that these TCR variants were nearly absent in Treg repertoires of healthy peptide-vaccinated mice (**Supplementary Fig. S3**). This suggested that growing tumor promotes either Th-to-Treg clonal plasticity or independent antigen-specific priming of convergent Th and Treg clones recognizing the same tumor neoantigens.

To analyze Th-to-Treg plasticity, we tracked identical nucleotide CDR3 sequence variants that revealed prominent clonal Th-Treg overlap in anti-CTLA4 treated but not in control mice. This may generally indicate that suppression of CTLA4-mediated Treg function elicits compensatory Treg programs in conventional helper T cells (41–43).

At the same time, TCRβ CDR3 variants specific to B16 melanoma neoantigenic peptides were not present among the clonotypes shared between Th and Treg nucleotide CDR3 repertoires.

We concluded that this presence of homologous yet distinct Th and Treg TCR sequence variants reflects independent priming of convergent Th and Treg clones specific to the same cancer neoantigens. This process can be mediated by the growing tumor’s influence on associated dendritic cells, thus supporting development of immunotherapeutic interventions that modulate dendritic cell behavior (44–48).

This dual activation leads to a competition within the tumor environment, where both immune suppression (via Tregs) and anti-tumor immunity (via effector CD4+ T cells) are simultaneously initiated. Tumor-induced DCs can secrete factors like indoleamine 2,3-dioxygenase (IDO), skewing the immune response towards Treg induction, which suppresses cytotoxic T cells and diminishes anti-tumor immunity (49–52). This balance or imbalance between effector T cells and Tregs, mediated by shared tumor antigenic specificity, is a key aspect of the immune regulation in cancer (45).

The convergence of TCR specificity in both effector and regulatory T cells might suggest a mechanism through which tumors manipulate immune responses to their advantage by tipping the scales in favor of immune suppression, thereby promoting tumor survival and growth. This concept supports development of therapeutic strategies that employ modulation of dendritic cells function or selectively target tumor antigen-specific Tregs for cancer immunotherapy.

In should be noted that our approach was based on vaccination with long 27-mer peptides and was inefficient in eliciting public CD8+ T cell clones. We suggest that use of 9-10-mer peptides or mRNA vaccination followed with a similar strategy of TCR repertoire profiling and comparative repertoire post-analysis may be efficient to identify CD8+ TCR motifs recognizing peptides of interest.

It should be also noted that melanoma B16F0 is evolutionarily younger than B16F10. B16F10 was obtained from B16F0 by 10 rounds of injecting tumor cells into the tail vein and selecting those neoplasms that engrafted/metastasized in the lungs. Here we tracked TCRs that are specific to peptides from the evolutionarily younger tumor variant, that may have arisen due to a new mutation in B16F10 and may be absent in original B16F0 line (53).

To conclude, here we:

1. Created a database of cross-mice shared (public) TCRβ CDR3 variants specific to 14 previously described B16 melanoma neoantigenic peptides in I-Ab pMHCII context of C57BL/6 mice, available to facilitate informed research and development in this preclinical model.
2. Report a wet lab and bioinformatics pipeline that is easy to reproduce to derive public Th TCR variants specific to other neoantigenic peptides, potentially also exploitable for CD8+ T cells with short peptides.
3. Among 14 neoantigenic B16 melanoma peptides, we identified most immunogenic in terms of relative abundance of responding Th cells in draining and non-draining lymph nodes in the orthotopic model, including peptides p5, p20, p25, p30, and p50.
4. Demonstrated that blocking-only anti-CTLA4 therapy promotes clonal expansions within Th but not within Treg subset, in both draining and non-draining lymph nodes.
5. Demonstrated that blocking-only anti-CTLA4 therapy promotes Th-to-Treg plasticity, observable in both draining and non-draining lymph nodes, but not specifically focussing on tumor antigens.
6. Demonstrated that, regardless of anti-CTLA4 therapy, B16 melanoma promotes clonally independent priming and expansion of Tregs that share sequence similarity with the Th TCRβ CDR3 variants specific to melanoma neoantigenic peptides.
7. Demonstrated that TCRβ CDR3 clusters derived from draining and non-draining lymph nodes’ Th repertoires are enlarged and enriched with TCRβ CDR3 variants specific to melanoma neoantigenic peptides upon blocking anti-CTLA4 treatment.

Altogether, we hope our work will facilitate informed investigation and understanding of the tumor environment and therapy developments based on the knowledge of TCRs characteristic for tumor-specific T cell clones.

## Supplementary

**Supplementary Figure S1.**
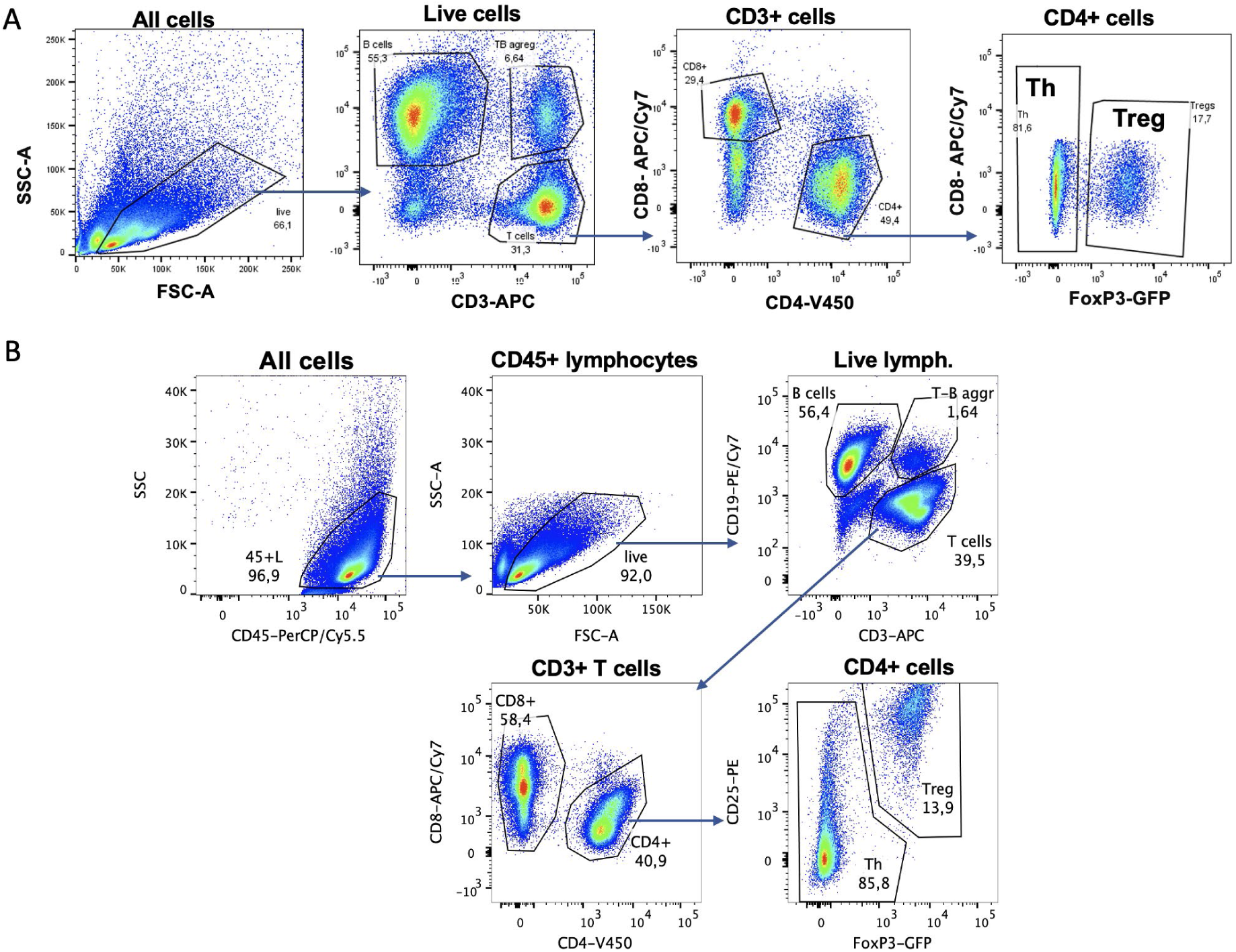
FACS sorting. **a**,**b**. Gating schemes for sorting cells from popliteal LNs (a), and inguinal LNs (b). In both cases CD8+, Th and Treg cells were sorted.

**Supplementary Fig. S2.**
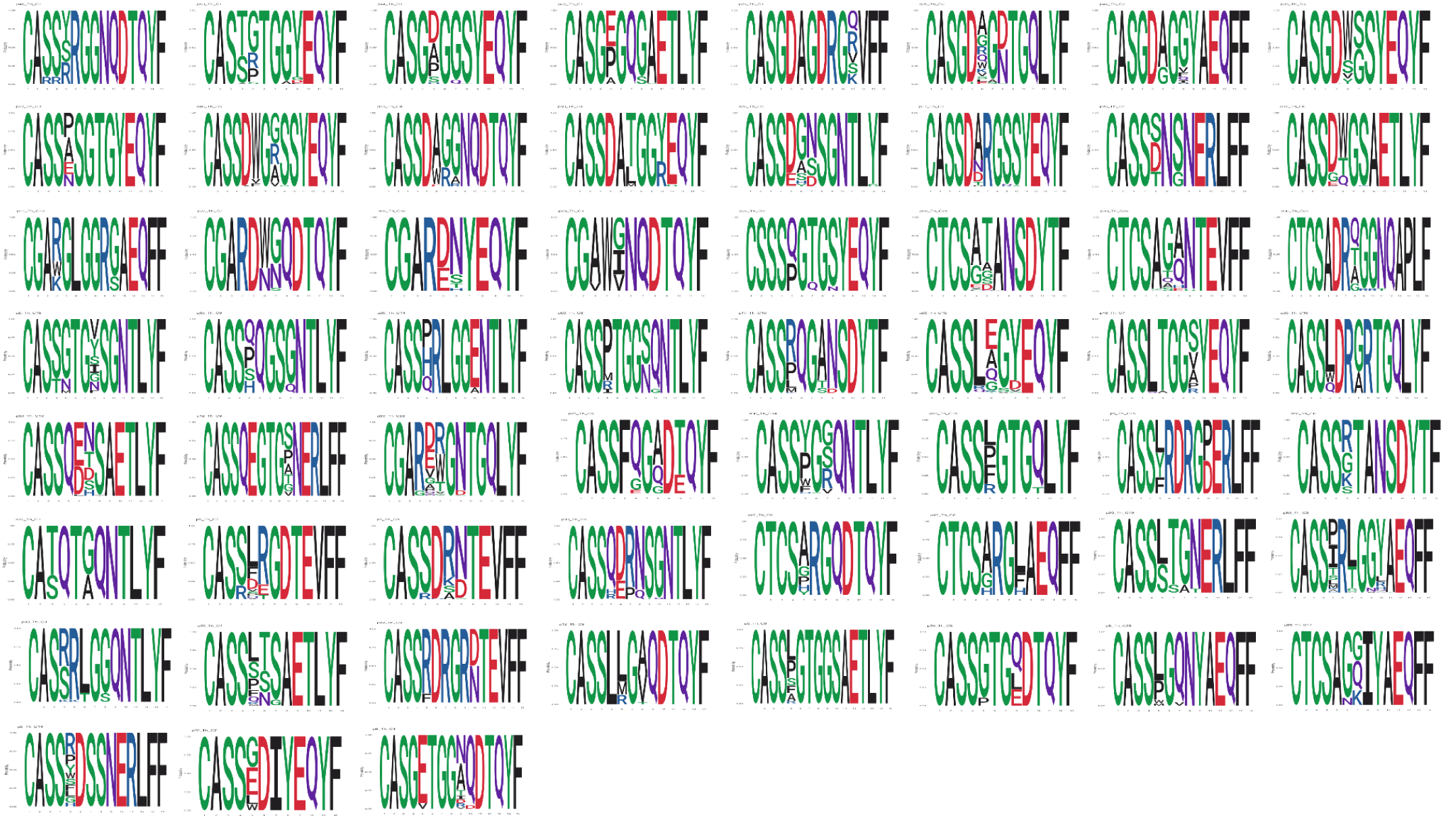
B16-specifc TCRβ CDR3 motifs. Logo images for identified B16 melanoma–specific TCRβ CDR3 clusters are shown.

**Supplementary Figure S3.**
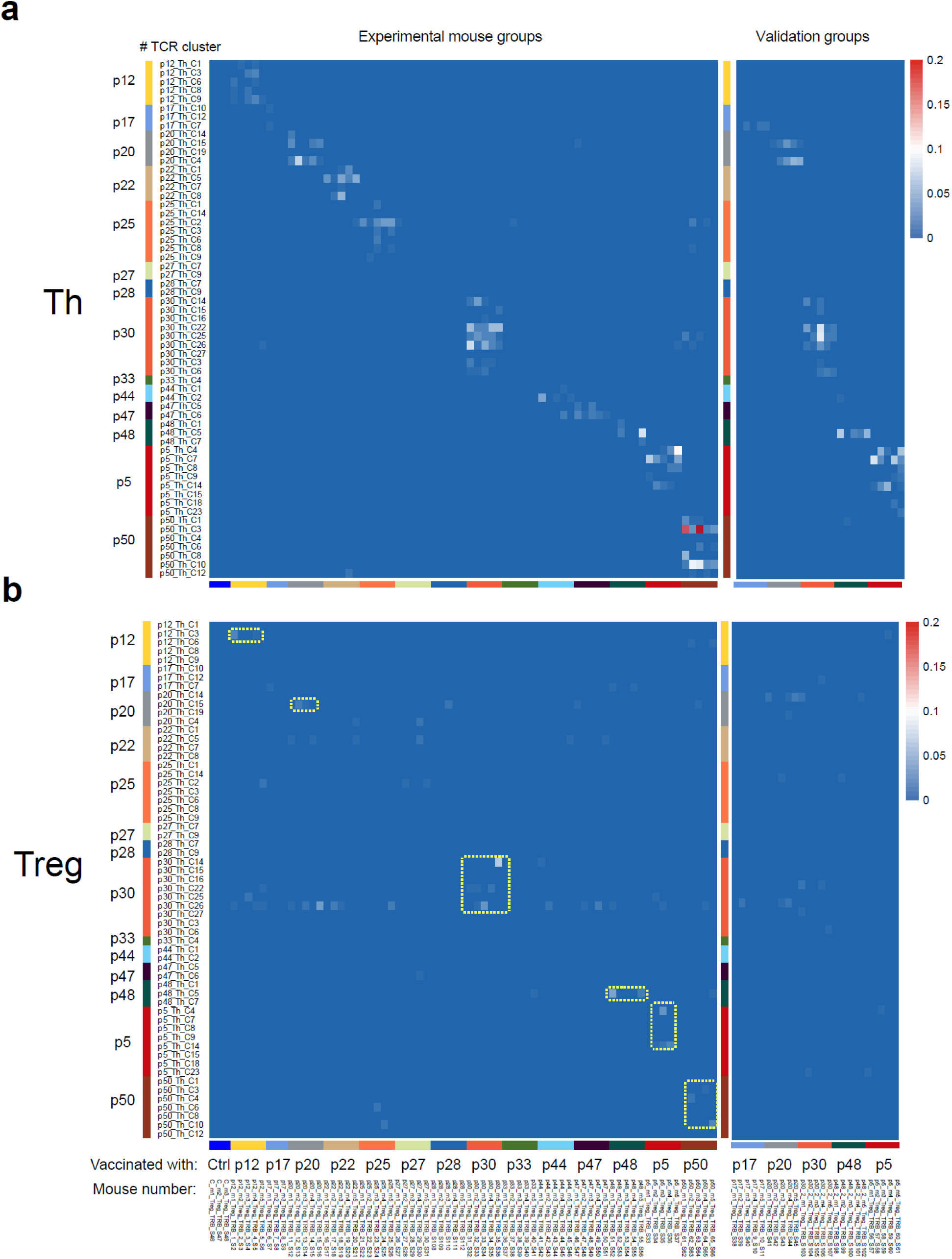
Distribution of identified B16F10-specific TCRβ CDR3 clusters with 1 amino acid mismatch allowed. Cumulative frequencies of all TCRβ CDR3s from each TCR cluster (rows) in each mice (columns) are shown for TCRβ CDR3 repertoires from discovery and validation mice groups. **a**. Th subset. **b**. Treg subset.

**Supplementary Figure S4.**
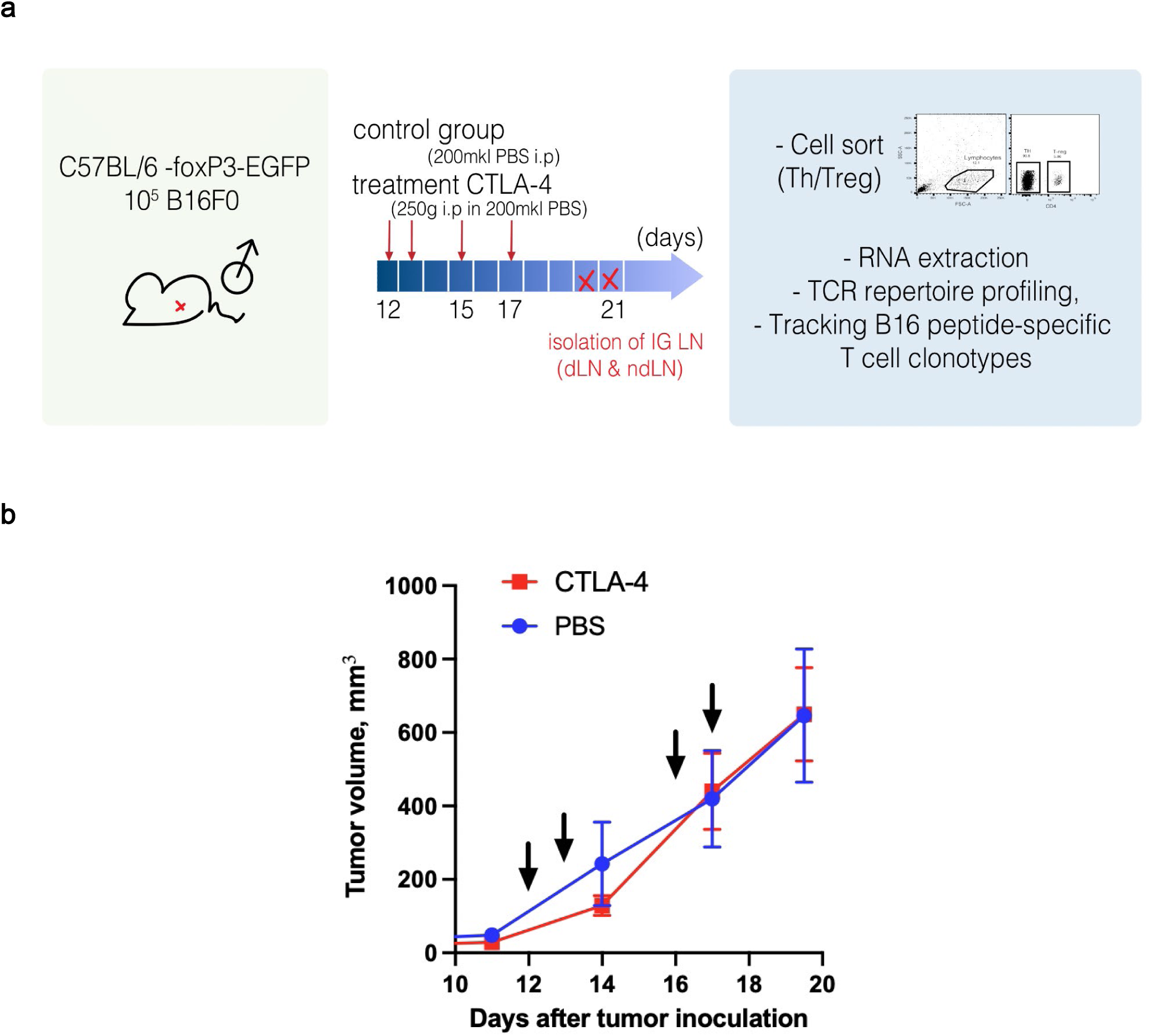
B16F0 melanoma orthotopic model. **a**. B16F0 melanoma orthotopic model, experimental scheme. **b**. Mean tumor volume for control mice (PBS, n=4) and anti-CTLA4-treated mice (n=7). Anti-CTLA4 antibodies were injected i.p. at days 12,13, and 16, 17 (indicated by arrows) and sacrificed on day 19, 20.

**Supplementary Figure S5.**
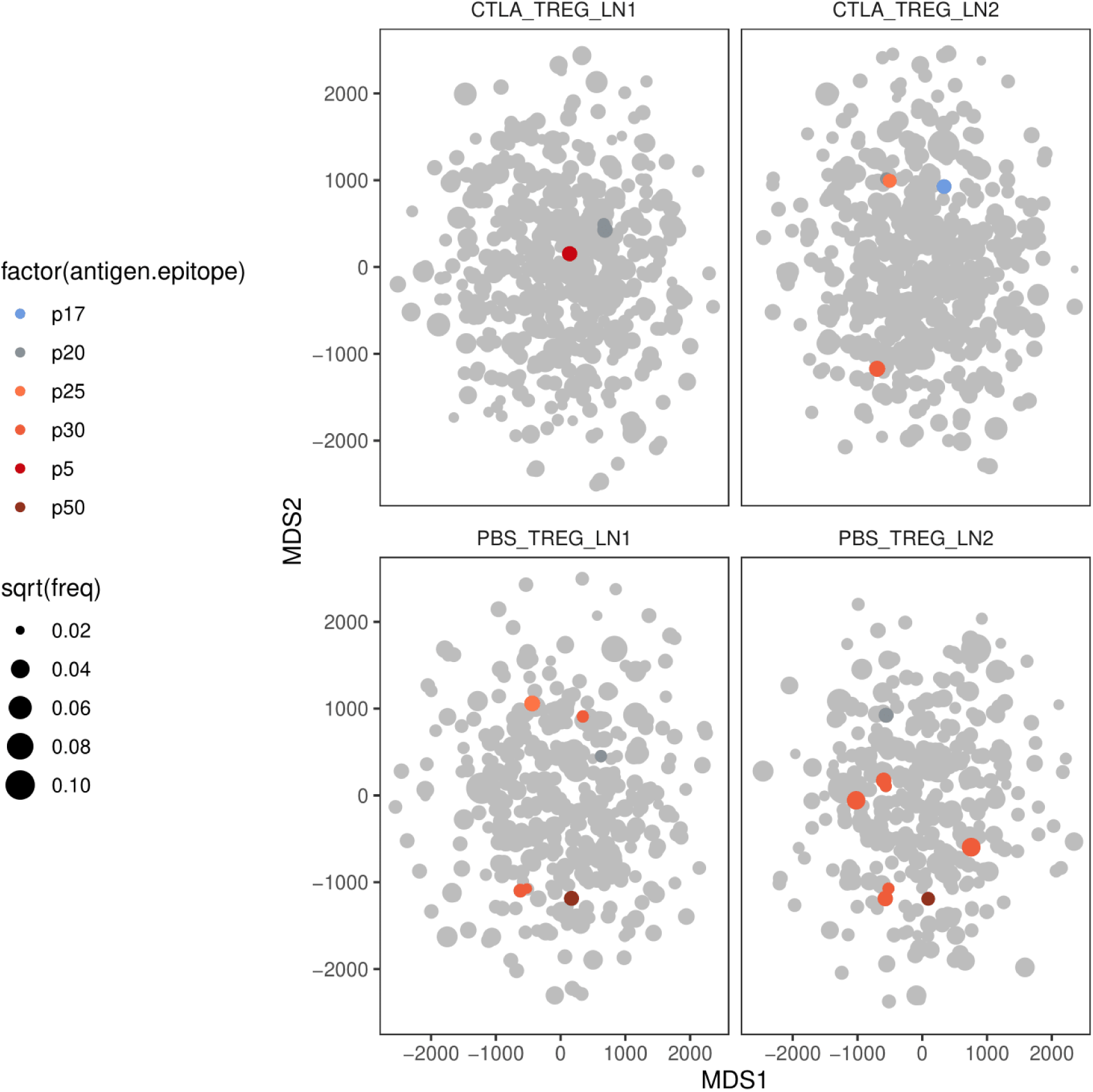
Treg TCRβ CDR3 clusters derived from tumor-bearing mice. Clusters were identified using TCRnet from the integrated top-500 clonotypes from Treg TCRβ CDR3 repertoires from dLNs and ndLNs of PBS and anti-CTLA4 treated mice. Each cluster is shown as a node. Clusters that match identified melanoma peptide-specific TCRβ CDR3 variants are colored.

